# Serotonin-dependent kinetics of bursts of feeding underlie a graded response to food availability

**DOI:** 10.1101/051086

**Authors:** Kyung Suk Lee, Shachar Iwanir, Ronen B Kopito, Monika Scholz, David Biron, Erel Levine

## Abstract

Animals integrate physiological and environmental signals to modulate their food uptake. Failure to regulate feeding may have devastating results, including obesity and diabetes, underscoring the importance of understanding its underlying mechanisms. The nematode *C. elegans*, whose food uptake consists of pumping bacteria from the environment into the gut, provides excellent opportunities for discovering principles of conserved regulatory mechanisms. Here we show that worms implement a graded feeding response to the concentration of environmental bacteria by modulating a commitment to bursts of fast pumping. Using long-term, high-resolution, longitudinal recordings of feeding dynamics under defined conditions, we find that the frequency and duration of pumping bursts increase and the duration of long pauses diminishes in environments richer in bacteria. The bioamine serotonin, a known feeding regulator in metazoa, is required for food-dependent induction of bursts as well as for maintaining their high rate of pumping through two distinct mechanisms. Following this phenotype quantitatively we identify the essential serotonergic neurons and the differential roles of distinct families of serotonin receptors. We propose that regulation of bursts is a conserved mechanism of behavior and motor control.

## Introduction

The regulation of food uptake is a critical mechanism with major physiological impacts. Aberrant feeding behaviors undermine weight regulation and are associated with a range of chronic diseases, while dietary restriction has implications on health and longevity ^1–6^. Simple model systems, like the nematode *C. elegans*, have proven invaluable to understanding the mechanisms of regulation of food uptake ^7,8^. Feeding of worms on bacteria is facilitated by the action of the pharynx, a neuromuscular pump that draws bacteria suspended in liquid into the gut from the surrounding environment and transports them to the intestine after concentrating and grinding ^7,9–13^. This process is conducted by a combination of two feeding motions, pumping and isthmus peristalsis. Pharyngeal pumping is a cycle of contraction and relaxation of pharyngeal muscles, in which bacterial food is taken from the environment, bacterial cells are trapped, and the surrounding liquid is expelled. Isthmus peristalsis is a peristaltic contraction of the muscles that carry the food within the pharynx ^7,12^. Pharyngeal pumping is therefore a primary indicator of food intake.

The two feeding motions are operated by the pharyngeal nervous system, one of the two independently functioning nervous systems of the worm, consisting of 20 neurons of 14 types ^14^. The MC and M3 neurons control pumping, while M4 is essential for isthumus peristalsis ^10,11^. Previous results suggest that pharyngeal pumping depends on feeding history and quality of food through a mechanism that involves the neurotransmitter serotonin (5-HT, 5-hydroxytryptamine) ^15–18^. Serotonin increases feeding rate of *Caenorhabditis elegans* ^19^ and has been suggested as a putative food signal that controls feeding of the animal ^20^. Evidence suggests that serotonin acts directly on MC and M3 neurons to facilitate fast pharyngeal muscle contraction–tion cycles ^16,21,22^.

Serotonin is a key modulator in the worm, involved in a variety of processes including foraging, mating, egg-laying, metabolism, chemosensation, aversive olfactory learning, and feeding ^20,23–30^. Endogenous synthesis of serotonin requires the tryptophan hydroxylase, which is expressed in all serotonergic neurons from the *tph-*1 gene ^14,25,31^. Two pairs of head serotonergic neurons, the pharyngeal secretory neurons (NSM) and the chemosensory neurons (ADF) ^18,20,32^, have been implicated in feeding, although their precise roles remain elusive ^32,33^. Additional complexity comes from the differential roles played by the different families of serotonin receptors. Three of the five known species of serotonin receptors, SER-1, SER-4, and SER-7, are expressed in pharyngeal neurons or muscles, suggesting a possible involvement in pharyngeal activity ^34–36^.

Here we hypothesize that worms modulate the dynamics of feeding in response to the availability of food in their environment, and we seek to characterize this response and trace its origins. Conventional feeding assays ^37^ are performed on dense bacterial lawns, which does not allow for fine control of food concentration. We therefore employed a custom microfluidic device ^38^ that enabled us to precisely control the concentration of available food and to monitor the dynamics of pharyngeal pumping at high resolution. Using these data, we show that feeding is characterized by bursts of fast, regular pumping, whose duration and frequency are correlated with the availability of food. In addition, feeding at low food densities is characterized by an abundance of long pauses. We implicate serotonin in promoting fast pumping and in suppressing long pauses and utilize these phenotypes to show that the serotonergic neurons NSM and HSN are together necessary and individually sufficient for fast pumping. This is the first time the HSN neurons, which are located in the middle of the worm body and send no processes into the pharynx, are implicated in the regulation of feeding. Two serotonin receptors, the 5-HT1 ortholog SER-1 and the 5-HT2 ortholog SER-4, are involved in food-dependent induction of fast pumping. While SER-1 receptors are involved in controlling the abundance of fast pumping, SER-4 are involved in maintaining the high pumping rate. Finally, we point out differences between induction of pumping by food and by exogenous serotonin, which relies on the 5-HT7 ortholog SER-7.

## Results

### Long-term automatic recording of pumping dynamics at defined food concentrations

Typical measurements of pharyngeal behavior involve manual scoring over short (30 - 60 sec) intervals ^37^, to estimate the average number of pumping events observed per unit of time, or the average pumping rate. To allow more detailed characterization of feeding dynamics, we performed long-term time-lapse imaging of the pharynx of multiple worms (**Fig. 1A** and **Movie S1**). For longitudinal measurements, worms were individually confined in a custom microfluidic device which permits maintenance of worms for longer than 24 hours with no detectable effect on their longevity, egg-laying, or pumping dynamics (**Supp. Fig. 1**) ^38^. Bacteria suspended in standard S-medium were constantly flown through the device, ensuring a continuous supply of bacteria at a fixed density. In what follows we indicate the densities of bacterial suspensions in units of optical density at 600nm (OD_600_).

**Figure 1.**
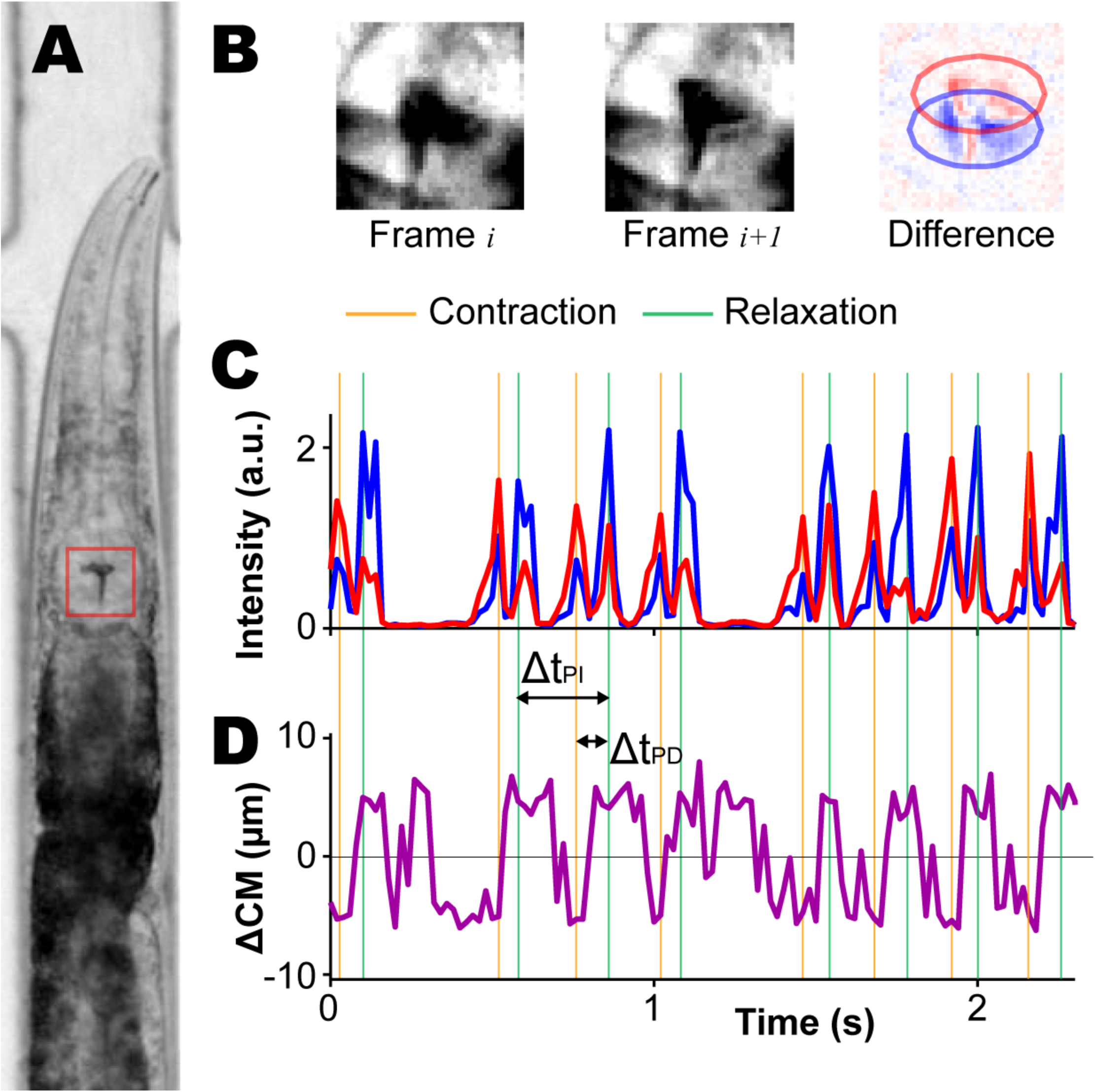
Automated detection of pumping dynamics. (A) Bright-field image of a worm confined in a custom microfluidic device for longitudinal recording of pumping dynamics. (B) Images of a small region surrounding the pharyngeal grinder (red box in (A)) at two consecutive frames and the difference between the two. Pixels that turn darker and pixels that become brighter are colored in red and blue, respectively. (C) Representative time courses of the total intensity of the red and blue marked pixels. Peaks in these measures are robust indicators of a pharyngeal motion, either contraction or relaxation. (D) Representative time courses of the displacement between the center of mass of the red-and blue-marked pixels. Positive values occur when the blue center of mass is posterior to the red one, indicating a relaxation motion.

In our experiments, worms were loaded to the device and fed high-density bacteria (OD_600_=3) for 3 hours before transitioning to the test media and beginning measurements. We cycled through the worms, taking a 5-30 minute time-lapse movie at a frequency of 50 frames per second for each worm in each imaging cycle. Our measurements lasted 2-3 hours, allowing us to revisit most worms multiple times.

The resulting movies were analyzed using a custom workflow that automatically detects pumping events by tracking the motion of the grinder. The high contrast between the grinder and its surroundings and its fast motion permit easy detection of pumping events by comparing consecutive images (**Fig. 1B**, **Movie S2** and Materials and Methods). The intensity of the difference between images is indicative of the speed of the grinder motion (**Fig. 1C**), while the displacement between changing regions provides information about the direction of the motion (**Fig. 1D**). A pumping event is identified as a reset of the grinder (after the previous event), inversion of the grinder plates, and relaxation of the muscle (**Fig. 1C,D**) ^12,13^.

For each event, we define Δ*t*_PD_ as the time between contraction and relaxation in a single pumping event (Pump Duration, PD), and Δ*t*_PI_ as the time interval between relaxation times of consecutive pumping events (Pumping Interval, PI; **Fig. 1C**). Since each contraction-relaxation cycle corresponds to a single muscle action potential _39,40_, the pulse duration Δ*t*_PD_ and the inter-pulse interval (IPI) Δ*t*_IPI_ = Δ*t*_PI_ – Δ*t*_PD_ are related to the duration of an action potential and the time the neuron membrane potential rebounds after repolarization, respectively.

### The average pumping rate, but not the duration of individual pumps, increases with food concentration

Food availability is presumed to be one of the key factors that regulate feeding. In worms, the presence of bacterial food on solid media stimulate pumping ^20^, and the mean feeding rate of worms grown in liquid culture increases with food concentrations ^19^, although worms grow slowly and show reduced fertility in liquid medium ^41,42^.

To characterize the regulation of feeding behavior in response to food availability, we collected and analyzed time-series data from multiple “wild-type” worms (the laboratory strain N2) at different food concentrations in the range from OD_600_=0 to OD_600_=8. To connect with previous results, we first estimated the *average* pumping rate at each density by counting the number of pumping events observed in each worm, dividing this number by the duration of that measurement, and averaging over all worms. We found that the average rate increases gradually with the concentration of supplied food from almost zero in clean media to a maximal average rate of 270 pumps/min in densities above OD_600_=3 (**Fig. 2A**).

**Figure 2.**
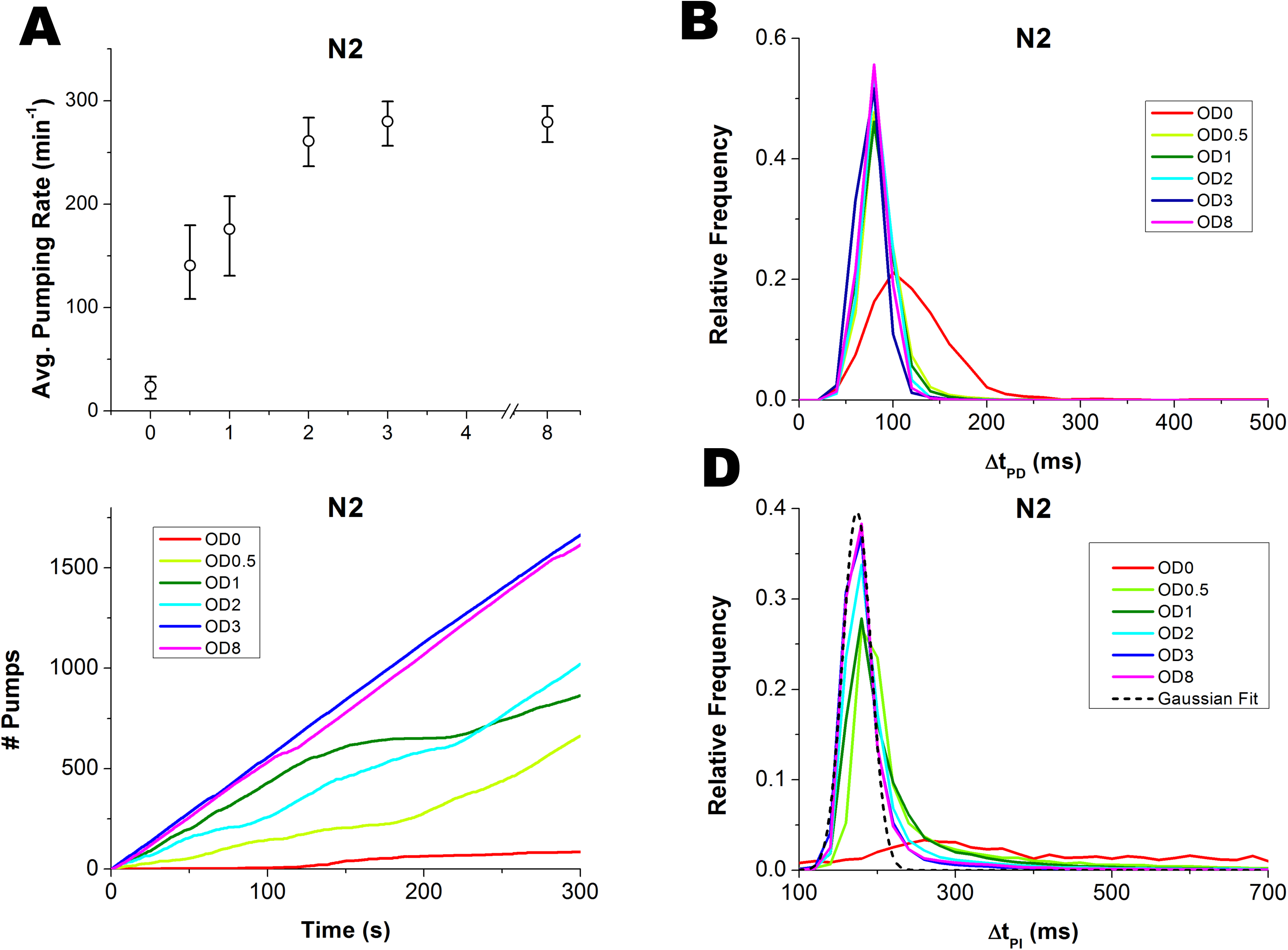
Pumping dynamics are modulated in response to food availability. (A) Average pumping rate as a function of food concentration. The rates are averaged over all worms and all recorded periods at each concentration. Error bars are 95% confidence intervals, calculated from an ensemble of all 10-minute subsets of the data. (B) The distributions of pump duration Δ*t*_PD_. (C) Representative time series of pumping by individual worms at different food concentrations. Plotted is the total numbers of pumps counted since the beginning of the experiment (defined as time=0). (D) Histogram of pulse intervals, Δ*t*_PI_. These histograms show a narrow peak and a heavy tail. One of these peaks was fit to a Gaussian distribution (black dashed line) centered at 174ms, with the standard deviation of 20ms (adjusted R^2^ = 0.985).

The duration of individual pumps was unaffected by the density of food. To see this, we measured the relative frequency (*i.e.*, the empirical probability) of pump durations Δ*t*_PD_ at different food concentrations. We found that these distributions were always sharply peaked around 80ms in all food concentrations, except in the complete absence of food (**Fig. 2B** and **Supp. Fig. 2A**). Thus, the duration of individual pumps is held fixed in fed worms, and does not contribute to the regulation of food uptake. The modulation of feeding dynamics is therefore captured in the way pumping events occur in time.

### A bursty pumping dynamics

An efficient way to represent the time-series of pumping by an individual animal is to graph the cumulative count of pumping events it performs over time (**Figs. 2C** and **Supp. Fig. 2B**). For each animal, the *instantaneous* pumping rate is the slope of a tangent line at any point on this graph, while the *average* pumping rate is the slope of a chord line of the same graph (**Supp. Fig. 2C**). At high food concentrations (above OD_600_=3), most worms pumped at a constant instantaneous rate, which was close to their average rate. In contrast, at lower food concentrations the pumping-count graphs alternated between periods of regular pumping at almost constant instantaneous rate (straight diagonal segments), and periods of much slower and irregular pumping (curved segments). In such cases, the average pumping rate fails to capture the true dynamics of feeding. This led us to hypothesize that pumping dynamics are not homogeneous in time. Rather, we propose that pumping occurs in bursts of fast, regular pumping, punctuated with periods of slower, irregular pumping.

Support for the idea of inhomogeneous pumping dynamics comes from the relative frequency of different pumping intervals (Δ*t*_PI_) observed at different food concentrations (**Fig. 2D**). These histograms were typically composed of a sharp peak around 174ms (corresponding to 6 pumps/second, or 345 pumps/min), and a heavy tail of much longer intervals, extending up to several seconds. The short-interval peak, resembling a Gaussian distribution with a standard deviation of 20ms, dominated the histograms at high food concentrations, consistent with the observation that at these densities, individual worms pump at a constant instantaneous rate (**Fig. 2C** and **Supp. Fig. 2B**). Previous electrophysiological recordings from the pharynx demonstrate that a pumping action potential and the subsequent rebound after repolarization take around 250ms ^13^, a timescale comparable to the pumping interval. This suggests that the pump intervals that contribute to the peak are as short as physiologically possible.

To help in classifying different pumping behaviors, **Fig. 3A** shows the same data as **Fig. 2D** in a different representation. For a given time *t*, we let *F*(*t*) be the fraction of time spent by the population of worms in pumping intervals (Δ*t*_PD_) that are longer than *t*. For example, F(600ms) = 10% means that on average a worm at the relevant food concentration spends 10% of its time in pumping intervals that are longer 600ms. In **Fig. 3A** we plot the measured values of *F* at different food densities as a function of the instantaneous rate *r=1/t*, rather than *t*, since this representation clearly suggests 3 classes of pumping dynamics: long pauses (corresponding to the sharp decline at small *r*); fast pumping (or short intervals, the steep increase at large *r*); and intermediate intervals, for which - curiously - the curve shows steady linear increase with the rate *r*. This feature allows us to unambiguously define the fraction of time spent in long pauses and the fraction of time spent in bursts of fast pumping respectively as the value of *F*(*r*) at the first point and the value 1 − *F*(*r*) at the last point of the linear segment in each curve in **Fig. 3A**. In almost all cases we studied, the former occurs around *r*=100 pumps/min (corresponding to intervals longer than 600ms) and the latter around *r*=250 pumps/min (or intervals shorter than 240ms). For simplicity, we therefore refer hereafter to intervals longer than 600ms as *long pauses* and to pumping at rate higher than 250 pumps/min *as fast pumping*, except when the range of the linear part of *F*(*r*) is significantly different.

**Figure 3.**
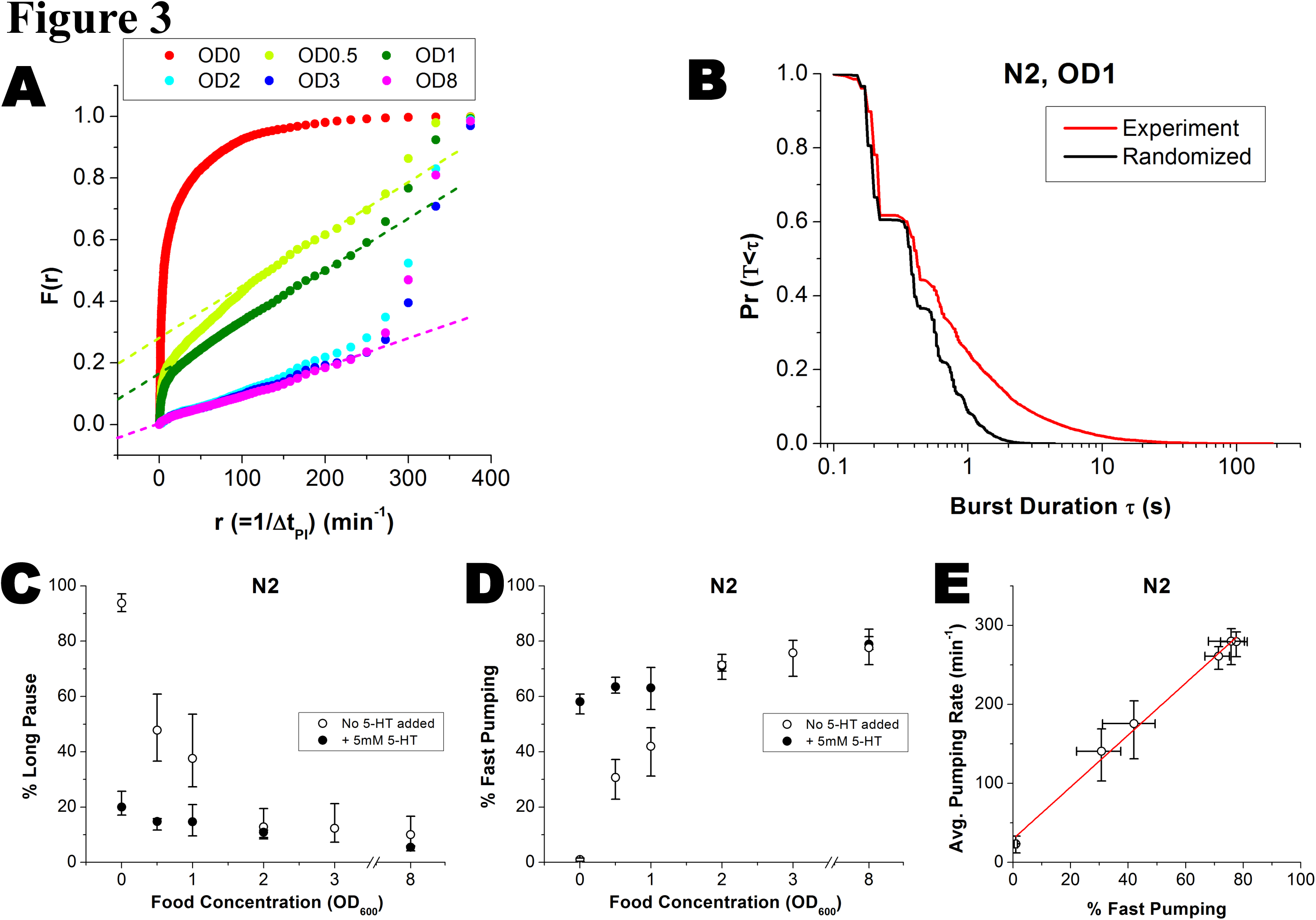
Bursts of fast pumping are a primary response to food availability. (A) *F*(*r*), the fraction of time spent in intervals longer than 1/*r*, at various food concentrations. Dashed lines for (OD_600_=0.5, 1, and 8) are the best-fit lines between 100min^−1^ and 250min^−1^. (B) Survival probability of the durations of fast pumping bursts at OD_600_=1 (black). Red curve is the burst durations calculated from the same pumping intervals in randomized order. (C and D) The average fraction of time spent by worms in long pauses (C) and fast pumping (D) at different food concentrations. Empty symbols, no added serotonin; filled symbols, in media supplemented with 5mM serotonin. Error bars as in Fig. 2. (E) Average pumping rates plotted against the fraction of fast pumping at different food concentrations. Red line is a linear fit to the data.

As exemplified in **Fig. 2C**, individual worms often display pumping dynamics of all three classes (**Supp. Fig. 3A**). The hypothesis that fast pumping appears in bursts suggests that short intervals (Δ*t*_PI_) are clustered in time, rather than spread randomly among intermediate intervals and long pauses. To test this hypothesis, we compared the observed statistics of burst duration (red curve of **Fig. 3B**) with what is expected if the duration of different intervals were uncorrelated (black curve of **Fig. 3B**). In support of our hypothesis, we observed that the duration of bursts was significantly longer than expected by chance (p-value < 10^−50^, Kolmogorov-Smirnov test).

### Modulating the commitment to fast pumping is the primary mechanism of adjusting feeding to food availability

We next sought to characterize the effect food availability has on the dynamics of pumping. The data in **Fig. 3A** suggests that the density of food affects the fraction of time spent in both fast pumping and long pauses. In worms not supplied with food, pumping dynamics were dominated by long pauses. These long pauses also take up a significant fraction of time in worms fed with low-concentration food, but the fraction of time spent in such pauses decreases rapidly with the increasing density of food (**Fig. 3C**). Conversely, the time spent in fast pumping increases gradually with the concentration of food (**Fig. 3D**). The relative time spent in both types of dynamics saturates around OD_600_=3.

In the presence of food, most pumping events occur during periods of fast pumping (**Fig. 2D**). We therefore reasoned that the modulation of the average pumping rate in response to food comes mostly from modulation of the fraction of time spent in fast pumping. Indeed, we find a very strong correlation (*R*^2^=0.99) between the average pumping rates at different food concentrations and the fraction of time spent in fast pumping (**Fig. 3E**). Variation in the fraction of time spent in bursts comes both from an increase in the average duration of individual bursts (**Supp. Fig. 3B**) and from an increase in their frequency (that is, a decrease in the time intervals between them, **Supp. Fig. 3C**).

### Serotonin production is required for fast pumping

The neurotransmitter serotonin is known to stimulate pharyngeal pumping by affecting MC neurons to increase the frequency of pharyngeal pumping and by decreasing the duration of the action potential in M3 neurons ^16,21,22,43^. In the absence of food, exogenous serotonin stimulates the activity of the MC neurons and induces pumping, mimicking the effect of food ^20^. As our results suggest that the relative frequencies of long pauses and bursts of fast pumping depend on food concentration, we asked if serotonin is involved in the mechanism behind this relation.

To address this question we first recorded the pumping dynamics in *tph-1* deletion mutants, which lack the tryptophan hydroxylase required for serotonin biosynthesis ^25^. The fraction of time spent in fast pumping was significantly suppressed in these worms at all food concentrations (**Fig. 4A**). In fact, fast pumping was under-represented in these worms, as seen by the absence of a high-rate steep increase in **Fig. 4B**. As a result, the average pumping rate in *tph-1* worms was significantly lower than in wild-type worms at all food densities (**Fig. 5A**).

**Figure 4.**
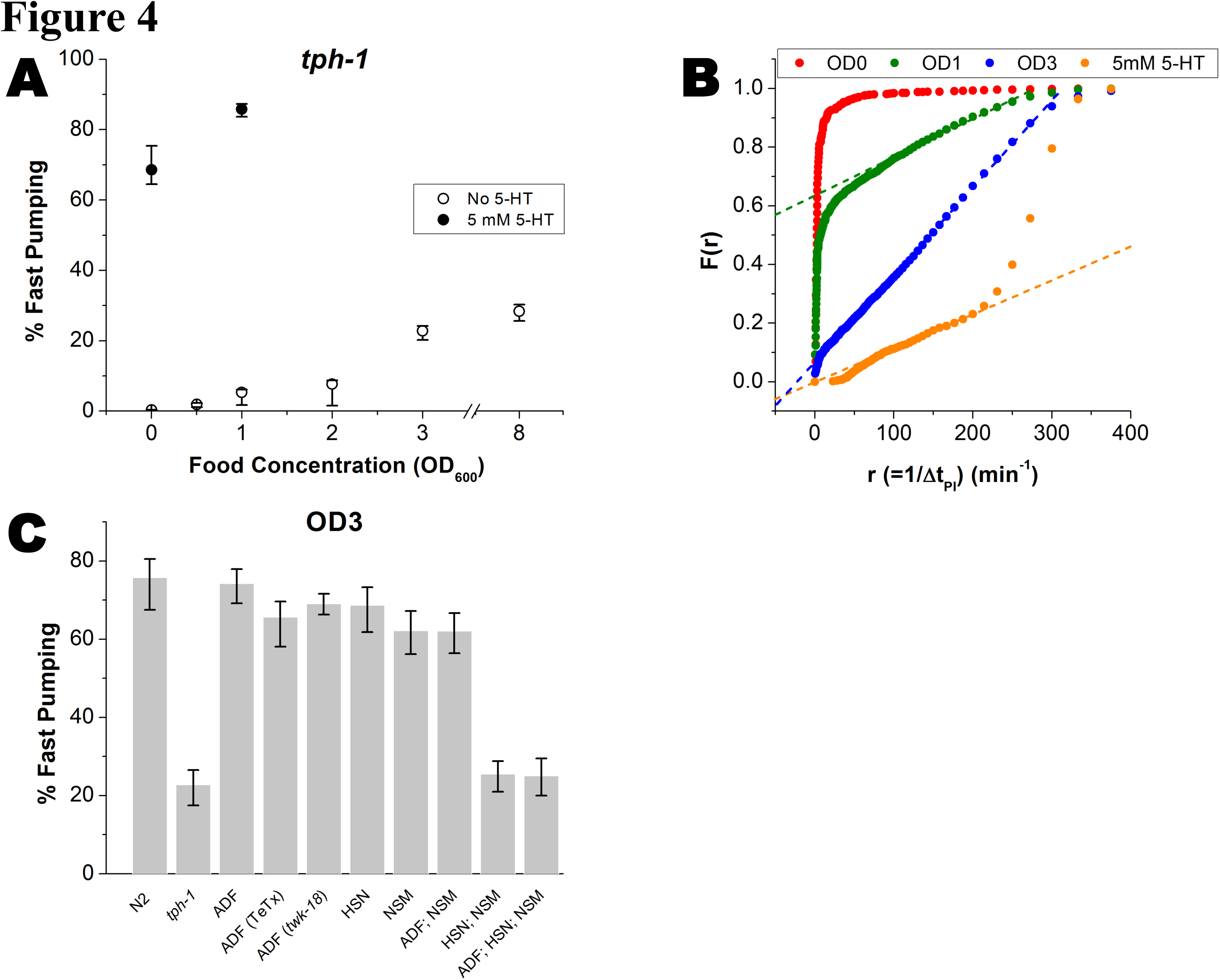
Serotonin is essential for induction of fast pumping. (A) The average fraction of time spent in fast pumping by *tph-1* worms at various food concentrations. Empty symbols, no added serotonin; filled symbols, in media supplemented with 5mM serotonin. (B) *F*(*r*), the fraction of time spent in intervals longer than 1/*r*, in various food concentrations with no serotonin added to the media (red, green, blue), and in 5mM serotonin in the absence of bacteria (orange). Dashed lines are the best-fit lines between 100min^−1^ and 250min^−1^. (C) The average fraction of time spent in fast pumping by mutants lacking serotonin production in different neurons, compared to wild-type (N2) worms and *tph-1* mutants at OD_600_=3. Error bars as in Fig. 2.

### Serotonin production in NSM and HSN is together necessary and individually sufficient for food-driven fast pumping

Two pairs of serotonergic neurons reside in the head of the worm, the sensory neurons ADF and the pharyngeal neurons NSM. Another pair of serotonergic motor neurons, HSN, is located in the middle of the worm body and known to control vulval and uterine muscles. While the ADF and NSM neurons were implicated in serotonergic modulation of feeding behavior ^18,20,32,44^, their role remains elusive. In addition, the HSN neurons are not known to be involved in regulation of pumping.

To examine the role of these serotonergic neurons in modulating feeding dynamics in response to food availability, we collected time-series of pumping from worms in which serotonin production was inhibited in subsets of these neurons. This was done using a Cre/Lox strategy, as previously described ^30^. Briefly, a single copy of the *tph-1* gene and its native promoter flanked by Lox signals was introduced into the genome of *tph-1* worms. The *cre* gene was then introduced into these worms using neuron-specific promoters (either individually or in groups), to excise the *tph-1* gene only in those neurons. Inhibition of serotonin production in all three pairs of neurons (NSM, ADF and HSN) simultaneously led to strong suppression of fast pumping (**Fig. 4C**) and the elimination of the high-rate steep increase (**Supp. Fig. 4A**) as observed in *tph-1* mutants (**Fig. 4B**), demonstrating the potency of the method.

In contrast, inhibition of serotonin production from each pair of neurons individually had a much less dramatic effect. Excision of the floxed *tph-1* either in the two ADF neurons or in the two HSN neurons had no noticeable effect on the dynamics of pumping. For ADF, this result is rather surprising, given the known role of ADF in regulation of feeding on agar surface ^32^ and for feeding response to familiar food ^18^. We therefore confirmed this result using two independent approaches. First, we inhibited synaptic release of serotonin from the ADF neurons by expressing the tetanus toxin (TeTx) from the ADF-specific *srh-142* promoter ^45^. These worms exhibited pumping dynamics indistinguishable from that of wild-type worms at both high and intermediate food concentrations (**Fig. 4C** and **Supp. Fig. 4B**). Similarly, inhibition of the ADF neurons by transgenic expression of an activated form of potassium channel *twk-18(cn110)* ^46^, showed no significant effect on pumping dynamics. Together, these results demonstrate that under these conditions, ADF neurons are not necessary for controlling the pumping dynamics, suggesting context-dependent functions of ADF.

Excision of floxed *tph*-1 in the NSM neurons alone or in both the NSM and ADF neurons, led to a small decrease in the fraction of time spent in fast pumping at high food concentration (**Fig. 4C**), but not at intermediate concentration (**Supp. Fig. 4B**) In contrast, simultaneous inhibition of serotonin production in NSM and HSN neurons led to a substantial decrease in fast pumping to the same level observed in *tph-1* worms and in worms where floxed *tph-1* was excised from all 3 pairs of neurons.

### The serotonin receptors SER-1 and SER-4 regulate different properties of food-driven fast pumping

Five serotonin receptors have been identified in *C. elegans* to date. Four are serotonin-activated G protein coupled receptors (SER-1, 4, 5 and 7) and one is a serotonin-gated Cl^−^ channel (MOD-1) ^36,47^. The different receptors have individual patterns of expression and different (sometimes antagonistic) roles in serotonin-dependent behaviors. Thus, the identities and functions of serotonin receptors involved in feeding behavior is a complex question. While the essential role of SER-7 in serotoninergic stimulation of pharyngeal pumping has received much attention ^16,18,43^, the excitatory roles of SER-1 ^16^ and SER-5 ^32^ as well as the inhibitory roles of SER-4 and MOD-1 ^18^ have also been discussed.

To investigate the roles of the different serotonin receptors in the modulation of fast pumping and long pauses in response to food, we considered worms carrying a null allele in each one of the known canonical serotonin receptors individually, or in a quintuple mutant carrying all five simultaneously. Unlike *tph-1* worms, which do not display bursts of fast pumping, all these strains showed consistent bursts in the presence of food. The average pumping rate, however, was significantly reduced in *ser-1* and *ser-4* mutants, as well as in worms carrying all five null alleles (**Fig. 5A**). In *ser-1* and in the quintuple mutants this was the result of similar suppression in the fraction of time spent in fast pumping (**Fig. 5B** and **Supp. Fig. 5B**). In contrast, *ser-4* worms did not show significant suppression in the time spent in bursts, but rather a reduction in the rate of their fast pumping. This was again determined by plotting *F*(*r*), the fraction of time spent in different pumping intervals (**Fig. 5C** and **Supp. Fig. 5A**), and identifying the fast rate as the rate of the final ascent, which occurs at 200min^−1^ in *ser-4* worms (cf. 250min^−1^ in wild-type). The reduction in the fast rate came from prolonged intervals between pumping events (Δ*t*_PI_) and not from change in the duration of individual events (Δ*t*_PD_) (**Supp. Fig. 5C**). Deletion of any one of the other three receptors — *ser-5, ser-7* or *mod-1* – showed no significant effect on pumping dynamics on food.

**Figure 5.**
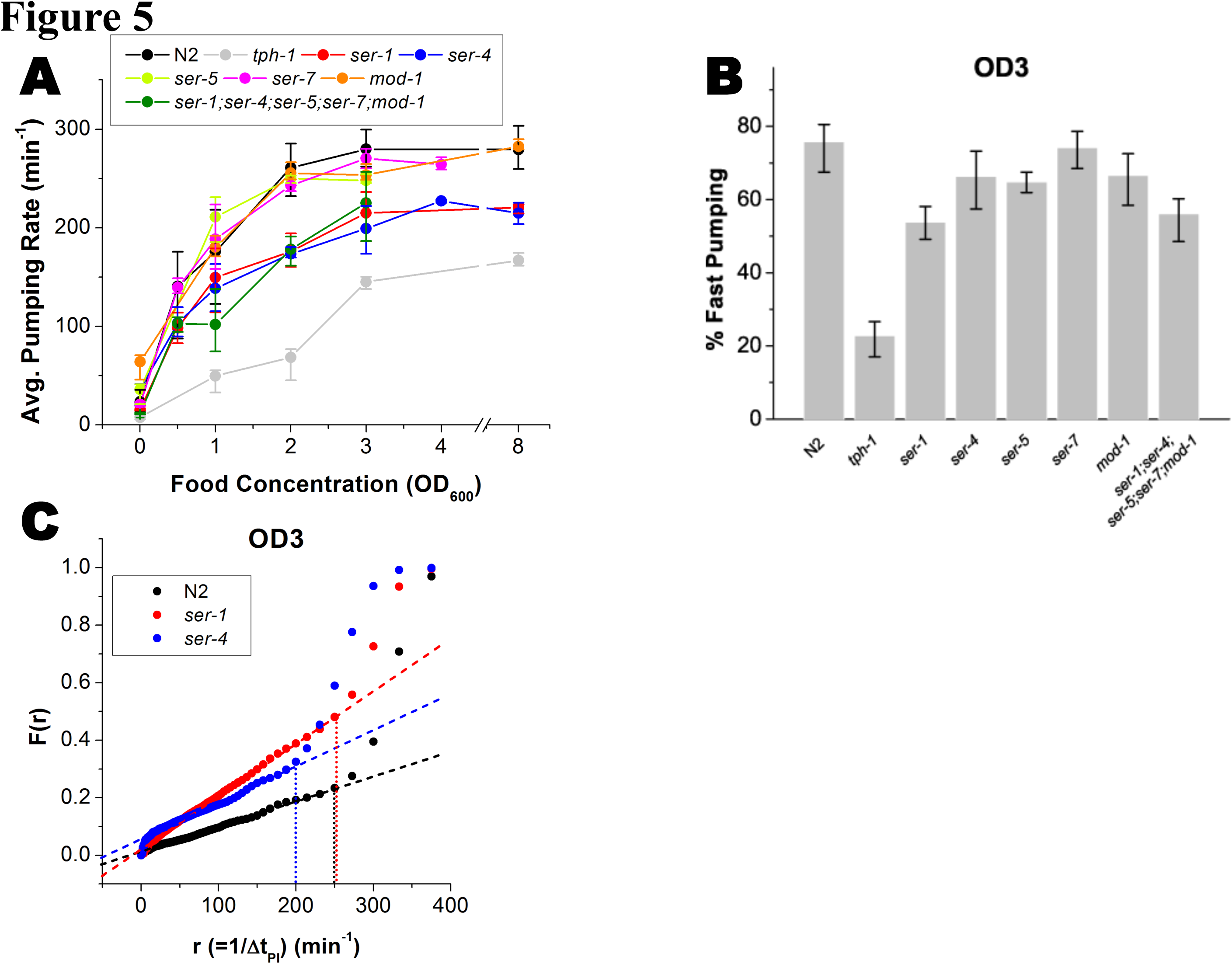
Distinct roles of serotonin receptors in food-driven pumping. (A) Average pumping rates of wild-type (N2) worms, *tph-1* mutants, and a range of serotonin receptor mutants at different food concentration. (B) The average fraction of time spent by these worms in fast pumping at OD_600_=3. Error bars as in Fig. 2. Here the threshold rate for fast pumping was taken to be 200min^−1^ for *ser-4* worms and 250min^−1^ otherwise (based on the next panel and on **Supp. Fig. 5A**). (C) *F*(*r*), the fraction of time spent in the intervals longer than 1/r, for wild-type worms and *ser-1* and *ser-4* mutants at OD_600_=3, with the best fit lines (dashed) between 100min^−1^ and 250min^−1^. The dotted vertical lines indicate the last point of the linear segment of *F*(*r*), taken as the threshold rate for fast pumping, demonstrating its slowdown in *ser-4* mutants.

### Exogenous supply of serotonin promotes fast pumping and suppresses long pauses

On agar plates, high concentration of serotonin activates overall feeding ^25,43^. To link this behavior to food-driven feeding, we recorded the dynamics of pumping in unfed wild-type worms supplied with serotonin at different concentrations (**Fig. 6A**). Mimicking the effect of food, higher concentrations of serotonin in the media led to higher average pumping rate. Also similar to food, increased concentrations of serotonin resulted in suppression of long pauses (**Fig. 6B**) and in an increase of the fraction of time spent in fast pumping (**Fig. 6C**). However, a clear difference was observed in the distribution of long pauses. In the presence of exogenous serotonin very long pauses, lasting several seconds, were effectively eliminated (**Supp. Fig. 6B, E**). Such long pauses were interrupted by a few pumping events, even when these did not form a significant burst (**Supp. Fig. 6C**). The dual effect of exogenous serotonin persists even in the presence of food (**Fig. 3C, D**). In addition, supplying exogenous serotonin allowed *tph-1* worms to exhibit fast pumping with dynamics similar to that of wild-type worms (**Fig. 4A, B**) ^43^.

**Figure 6.**
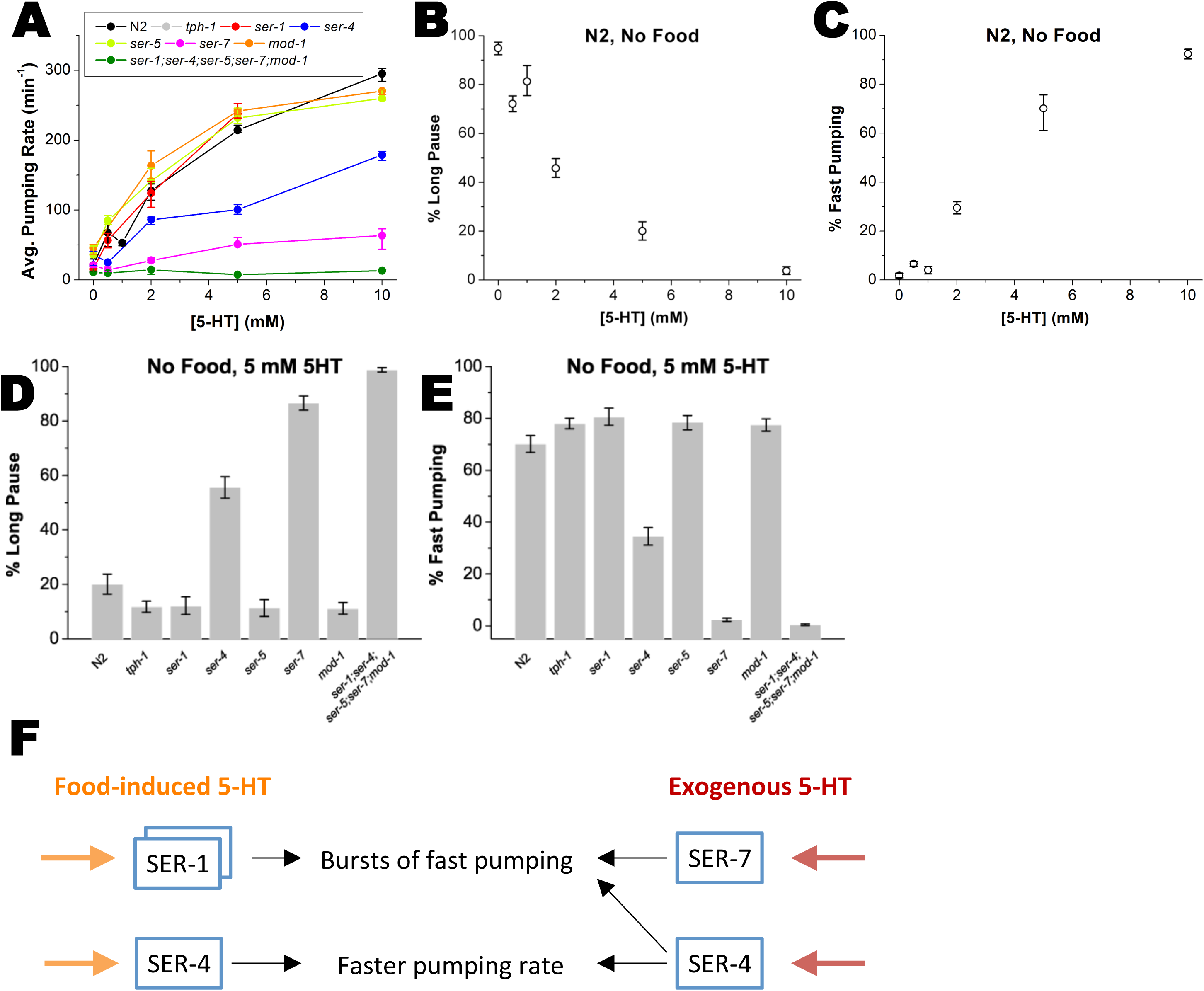
Distinct roles of serotonin receptors in pumping stimulated by exogenous serotonin. (A) Average pumping rate of wild-type (N2) worms, *tph-1* mutants, and a range of serotonin receptor mutants at different concentrations of serotonin (5-HT) in bacteria-free media. (B, C) The average fraction of time spent in long pauses (B) and fast pumping (C) by wild-type (N2) worms as a function of the concentration of serotonin in bacteria-free media. (D, E) The average fraction of time spent in long pauses (D) and fast pumping (E) by wild-type (N2) worms, *tph-1* mutants, and a range of serotonin receptor mutants in bacteria-free media supplemented with 5mM serotonin. Here the threshold rate for fast pumping was taken to be 160min^−1^ for *ser-4* worms and 200min^−1^ otherwise (based on **Figs. S6A** and **S6F**). Error bars as in Fig. 2. (F) Summary of the distinct roles of serotonin receptors. In food-driven pumping, SER-1 is involved in promoting pumping bursts, while SER-4 is required for maintaining their fast pumping rate. The stimulation of pumping by exogenous serotonin requires the SER-7 and SER-4 receptors. In this case, SER-4 is involved not only in establishing the rate of the bursts, but also in their induction.

Induction of feeding by serotonin is known to depend on the serotonin receptor SER-7 [38]. Indeed, even in the presence of high concentrations of serotonin (5mM), *ser-7* worms spent a considerable fraction of their time in long pauses (**Fig. 6D**) and displayed no fast pumping (**Fig. 6E**). A similar but weaker phenotype was observed in *ser-4* worms for both effects. As in food-driven pumping, the rate of fast pumping by these animals was slower than that of wild-type worms (**Supp. Fig. 6F**). Worms lacking all five canonical receptors showed the stronger phenotype, showing almost no response to exogenous serotonin and spending most of their time in long pauses.

## Discussion

The energy needs of a living organism must be fulfilled and balanced despite large fluctuations in nutrient availability and energy expenditure. Energy uptake is therefore regulated in response to the internal state of the body and to environmental conditions. Simple model organisms, like *C. elegans*, are particularly useful for elucidating underlying regulatory mechanisms. The advance in microfluidics and image processing techniques offers a unique opportunity for a longitudinal study of the regulatory response of living animals to precisely defined environments.

In this work, we showed that the rate of pharyngeal pumping in the worm is continuously matched with the concentration of bacterial food in the environment. We found that the time series of pumping in individual worms occurs in bursts of fast pumping. During these bursts, worms display fast, regular pumping, at a rate similar to the maximal rate allowed by the underlying neuronal physiology ^13^. The response in average feeding rate to food availability is controlled by modulating the fraction of time spent in fast pumping, rather than by continuously tuning the instantaneous pumping rate. Slow and irregular pumping has been previously described and linked to the effect of mutations that affect the MC neurons or the effect of starvation ^11,13,21,48^. Our data suggest, however, that slow pumping also occurs in healthy animals even in the presence of food as part of regulation of the overall feeding rate.

The biogenic amine serotonin (5-hydroxytryptamine; 5-HT) plays a pivotal role in energy homeostasis, *e.g.* by coordinating the perception of nutrient availability and feeding. Worms that cannot synthesize serotonin are capable of fast pumping, but fail to generate bursts. This result is consistent with the seminal work of Hobson *et al* ^16^, which showed how worms fail to maintain fast pumping over time when feeding on lawns of bacteria on agar plates.

We demonstrated that serotonin production in the pharyngeal motor neurons NSM and in the mid-body motor neuron HSN is necessary and sufficient for fast pumping in response to food. The involvement of NSM is consistent with previous reports that implicate the serotonin-controlled MC neurons in regular pumping ^11,13,16^ and complements studies of feeding under high food concentration conditions ^32,33^. On the other hand, the role of HSN in the regulation of feeding is unexpected, due to their distal location from the pharynx and their known roles ^49^. Recent findings, however, demonstrate a synergistic role for HSN and NSM in promoting a locomotion behavioral state known as dwelling, where worms inside a bacteria lawn move slowly and turn frequently to explore locally. This behavior relies on serotonin production in *both* the NSM and HSN neurons, but not in ADF ^26,50^.

That serotonin production in HSN alone is sufficient to elicit food-dependent pumping dynamics suggests two possible models. First, it is possible that serotonin produced by HSN is being taken up by the NSM neurons (through the reuptake receptor MOD-5, which is expressed in the NSM neurons) and is used by NSM to transmit food-dependent information to its post-synaptic cells. In this model, serotonin from HSN can be used by the NSM as an alternative source of this neurotransmitter in the absence of its *tph-1*. An alternative model suggests that humoral serotonin from HSN promotes fast pumping directly, without the need for NSM, supported by the fact that laser ablation of the NSM neurons had no noticeable effect on pumping dynamics in worms feeding on a plate ^11,18^. In this model, HSN activity depends on food availability, consistent with the fact that egg-laying rate, which is directly regulated by HSN, is significantly higher in the presence of abundant food than in its absence ^20^.

The serotonergic sensory neurons ADF have been previously implicated in pumping on plates and regulation of feeding in response to familiar food ^18,32^. It was therefore surprising that ADF had no role in modulating pumping dynamics in our experiments, and we verified these results using three independent approaches: excision of *tph-1* exclusively in the ADF neurons, inhibition of synaptic release of serotonin from ADF by expressing the tetanus toxin (TeTx), and inhibition of the ADF neurons by transgenic expression of an activated form of potassium channel TWK-18. In addition, previous studies of the roles played by serotonergic neurons in food-associated stimulation of pharyngeal pumping led to contradicting results supporting either ADF or NSM ^32,33^.

Previous studies have linked ADF with stress response ^28,51,52^ and feeding in response to change of food quality ^18^ The role of ADF in feeding regulation has been demonstrated for worms on a plate, where worms move freely in and out of the bacteria-covered region ^32^. In contrast, in our experiments, well-fed worms were assayed in stationary environments, where the animals are kept at constant food concentration for hours. Thus, it is possible that the roles played by the sensory neurons ADF are related with sensing fluctuations in the environment and are suppressed in our experiments. Consistently, we did not observe a clear phenotype in the mutant lacking *ser-5*, which has been associated with ADF-dependent feeding ^32^.

Five serotonin receptors have been previously identified in the worm. These receptors exhibit differential expression patterns and have distinct, sometimes antagonistic roles in different behaviors. With this in mind, we characterized the roles of these receptors in modulation of long pauses and fast pumping in response to food. Both *ser-1* and *ser-4* worms showed reduced pumping in the presence of food, but for very different reasons: *ser-1* worms spent less time in fast pumping than wild-type worms, while the rate of fast pumping of *ser-4* worms was slower. As *ser-4* seems to have no effect on the duration of individual pumps (**Supp. Fig. 5C**), *ser-4* is likely to be involved in invoking the contraction, rather than relaxation, of pharyngeal muscles. Together, our results suggest that SER-1 receptors are involved in promoting bursts of fast pumping, while SER-4 receptors are involved in establishing their fast rate. In further support of a role for SER-4 in regulation of feeding, it has previously been shown that serotonin-induced pumping is more suppressed in *ser-7; ser-4* double mutants than in the *ser-7* worms ^16^, and *ser-4* larvae pump significantly slower than wild-type larvae at the L1 stage when supplied with serotonin ^18^.

Activation of pumping by exogenous serotonin is markedly different from the dynamics induced by food. In the presence of serotonin at constant concentration, we could not observe long pauses that extend beyond a few seconds **(Fig S6E)**, while such pauses and longer were observed in feeding worms in all food concentration (**Supp. Fig. 6B**). An increased concentration of food induces an increase in average pumping rate and in fast pumping, with complete correlation between the two (**Fig. 3E**). While serotonin induces both fast pumping and an increase in the average pumping rate, the two were only weakly correlated (**Supp. Fig. 6D**). Also, the rate of fast pumping induced by exogenous serotonin was slightly lower than that of food-induced pumping (**Supp. Fig. 6A**). Finally, as noted before, serotonin-induced activation of pumping requires the SER-7 receptors, whose absence has no significant effect on food-driven pumping. Together, our results suggest a model, in which food-driven fast pumping involves the SER-1 receptors, stimulated pumping by exogenous serotonin involves the SER-7 receptors, and both require the SER-4 receptors for maintaining fast pumping close to the physiological limit (**Fig. 6F**). Interestingly, previous findings show that feeding on bacterial lawns on agar plates, *ser-1* and *ser-*7 mutants pump at an average rate similar to that of wild-type animals, but that their pumping is somewhat more irregular, while *ser-1; ser-7* double mutants fail to maintain fast pumping and display reduced pumping rate ^16^. This suggests a possible role for SER-7 receptors in food-driven bursts of pumping, which in our assay is insignificant in the presence of *ser-1*.

Serotonin facilitates a sharp response to food availability; in its absence, the feeding rate increases gradually with food concentration (*tph-1*, **Fig. 5A**). At OD_600_=1, for example, wild-type worms pump at 65% of their maximal rate, while *tph-1* worms pump at only a quarter of that rate. This suggests that the gain in fitness achieved by uptake of low concentration food exceeds the energetic costs of efficient pumping. In support of this idea, we recently found that stress-related activation by the Insulin/IGF-1 pathway in response to food limitation occurs only at food densities below OD_600_=1 ^53^. We also note that the average pumping rates of *tph-1* mutants were 28% and 52% of the wild-type rates at OD_600_=1 and 3, respectively, while a previous study found it to be 80% of the wild-type rate in worms feeding on lawns of bacteria on agar plates ^32^ (**Supp. Fig. 5D**). Taken together, these results suggest that serotonin regulates feeding rates in a context dependent manner.

Coupling between microfluidics and automated imaging enabled us to collect long-term longitudinal data from a large number of worms at high temporal frequency and in precisely defined environments. With these data we were able to identify the multi-faceted nature of the pumping dynamics, quantitatively link attributes of the environment and of the animal’s behavior, and characterize the distinct roles of serotonergic neurons and serotonin receptors.

In rodents, feeding is organized by bursts of licking, and the increase in feeding following starvation is mainly due to shortening of the interval between bursts (the OFF mode) ^54,55^. A recent study based on automated imaging of feeding behavior in *Drosophila* demonstrated a similar behavior in flies ^56^. Increase in feeding after short-term starvation was achieved by a decrease in the interval between bursts, while a long-term starvation resulted in an increase of the burst length. It is therefore possible that modulation of bursty feeding dynamics is a conserved strategy. In addition, the role of serotonin as a neuromodulator of feeding is conserved from invertebrates to mammals. The approach and tools described in this work, along with the relative simplicity of the worm’s neuronal circuitry and anatomy, open the way to further investigate the dynamics of feeding and the coupling between physiological and molecular processes, neuronal circuitry, and metabolism.

## Methods

### Strains

All strains were maintained on standard nematode growth medium (NGM) plates seeded with *E. coli* strain OP50 ^57^ at 15 °C. Adult hermaphrodites were used in all assays. The N2 Bristol strain was used as wild type animals, in addition to the following strains: MT15434 *tph-1(mg280)II*, INV33006 *Ex [tph-1p::TeTx::mcherry unc-122p::gfp]*, INV30001 *Ex [srh-142p::TeTx::mcherry unc-122p::gfp]*, ZC1890 *mgIS71V; yxEx960 [srh-142p::twk-18(cn110)::mcherry; unc-122p::dsred]* ^46^, CX13571 *tph**-**1(mg280)II; kySi56 IV; kyEx4077[srh**-**142p::nCre]*, CX13572 *tph-1(mg280)II; kySi56 IV; kyEx4057[ceh-2p::nCre]*, CX13576 *tph**-**1(mg280)II; kySi56 IV; kyEx4107[egl-6p::nCre]*, CX15658 *tph-1(mg280)II; kySi56 IV; kyEx5262 [ceh-2p::nCre, egl-6p::nCre]* ^30^, DA1814 *ser-1(ok345)X*, AQ866 *ser-4(ok512)III*, RB2277 *ser-5(ok3087)I*, DA2100 *ser-7(tm1325)X*, and RWK3 *ser-5*;*ser-4*;*mod-1*;*ser-7 ser-1* ^47^, ERL76 *tph-1(mg280)II; kySi56 IV; opyEx18 [ceh-2p::nCre, srh-142p::nCre, myo-3::mCherry]*, ERL82 *tph**-**1(mg280)II; kySi56 IV; opyEx19 [ceh**-**2p::nCre, srh-142p::nCre, egl-6p::nCre, myo-3::mCherry]*.

### Imaging pharyngeal pumping dynamics in a microfluidic device

For feeding experiments, an overnight culture of *E. coli* OP50 grown in LB media was centrifuged and resuspended at the required concentration in S-medium ^58^. The bacterial solution was filtered through a 5μm syringe filter to remove big aggregates, which could block the flow in the microfluidic device.

Age-synchronized worms were obtained by letting 15~20 gravid adults, incubated at 25 °C for at least a day, lay eggs on a plate for an hour. Adult worms were then removed, and the plates were incubated at 25 °C for 56~60 hours. Worms were mixed into a bacterial solution and loaded into the microfluidic devices (**Supp. Fig. 1**) as described previously ^38^.

Bacterial solution was injected into the device at the rate of 10μL/min. To ensure that bacteria do not clog the device, we applied a one-minute pulse of 150μL/min up to 3 times/hour. Data acquisition started 3 hours after feeding the animals at constant food concentration of interest. For the titration experiments (**Fig. 5A** and **Fig. 5B**) of strains other than N2, the same set of worms was monitored at multiple conditions, as we decrease the concentrations of food and exogenous serotonin, respectively, step by step, with 30-minutes wait time at each step.

Imaging was done using a Zeiss Axio Observer.Z1 microscope with N-Achroplan 10x objective (NA 0.25), and a Hamamatsu ORCA Flash4.0 camera. Every worm was imaged for 5 minutes at 50 frames/sec at spatial resolution of 0.65μm/pixel. When required, representative worms were recorded for 30 minutes.

### Image analysis

To analyze time-lapse images, the position of the grinder was identified in each frame. The difference in the brightness of pixels between every pair of consecutive frames was computed to detect contraction of pharyngeal muscles and subsequent relaxation (**Movie S2**). We identified the center of mass (CM) of pixels that become darker (marked in red in **Fig. 1B and C**) and the one of pixels that become brighter (blue). We then calculated the total difference in pixel brightness in an ellipse of constant area around each (**Fig. 1C**). Peaks in this measure are clear indicators of a rapid change in the pharynx position, either contraction or relaxation. To differentiate the two, we used the fact that the center of mass of the red pixels was posterior (ΔCM < 0) to that of the blue pixels during contraction and anterior (ΔCM > 0) to it during relaxation (**Fig. 1D**). Comparison of the total difference in brightness with the image background provides a confidence measure. Low confidence events were inspected manually and were either validated or discarded from further analysis.

## Acknowledgments

We thank Carlos Riberio and Yun Zhang for discussions and Richard Komuniecki, Mark Alkema, Yun Zhang and Cori Bargmann for reagents. This research was supported by the National Science Foundation through grants PHY-1205494 (EL) and IOS-1256989 (DB).

---

**Supplementary Figure 1. A worm in the microfluidic device**. Worms in the device feed on a bacterial suspension that is continuously flown through the chamber. The pillars restrain the worms from escaping while providing enough space for egg laying. Eggs are collected in an egg-collection area.

**Supplementary Figure 2. Pumping dynamics depend on food concentrations**. (A) Food concentration dependence of average pump duration time (Δ*t*_PD_). The bar graph shows average pump duration time (Δ*t*_PD_) with standard errors at respective food concentrations. (B) The number of pumps observed since the beginning of the experiment (time 0) for 3 different worms at each of 3 densities (OD_600_=0.3, 1, and 3), demonstrating the reproducibility of the assay. (C) Comparison of measured pumping counts (dotted curves) with their chord line (i.e. the lines that connect the two end points of the measured curves). The slopes of these chord lines are the average pumping rates

**Supplementary Figure 3. Bursts of fast pumping are longer and more frequent at higher food densities**. (A) A representative time series of pharyngeal pumping of a wild-type (N2) worm at OD_600_=1. Each pumping event is indicated by its instantaneous rate, i.e. the inverse of the time interval between that pump and the next. The time series shows long pauses, fast pumping, and pumping at intermediate rates. Blue and red dashed lines indicate the thresholds for fast pumping (250min^−1^) and long pauses (100min^−1^), respectively. (B, C) Survival probability of burst duration (B) and the time between bursts (C) at various food concentrations.

**Supplementary Figure 4. Serotonin production in HSN and NSM neurons is together necessary and individually sufficient for fast pumping**,. (A) *F*(*r*), the fraction of time spent in intervals longer than 1/*r*, at OD_600_=3. ADF(TeTx): strain in which the tetanus toxin is expressed from an ADF-specific promoter; *ADF(twk-18):* strain in which the activated form of potassium channel *twk-18* is expressed from a ADF-specific promoter; All others: strains in which *tph-1* has been removed specifically in the indicated neuron(s) using the Cre/Lox approach. (B) The average fraction of time spent in fast pumping by the same worms at OD_600_=1, compared to wild-type (N2) worms and *tph-1* mutants. Error bars as in Fig. 2.

**Supplementary Figure 5**. (A) *F*(*r*), the fraction of time spent in intervals longer than 1/r by serotonin receptor mutants, at OD_600_=3. (B) The average fraction of time spent by these worms in fast pumping at OD_600_=2. Error bars and thresholds as in Fig. 5. (C) Histograms of pump duration (Δ*t*_PD_) in *ser-4* at various food concentrations (colored curves), compared to wild-type (N2) at OD_600_=3 (black, dashed). (D) Ratio between the average pumping rate of *tph-1* worms and that of N2 worms at various food concentrations. Included for comparison is the same ratio for worms feeding on a lawn of bacteria on agar plates, as measured in ^32^.

**Supplementary Figure 6. Stimulated pumping by exogenous serotonin**. (A) *F*(*r*), the fraction of time spent by wild-type (N2) worms in intervals longer than 1/*r* in bacteria-free media supplemented with exogenous serotonin. For comparison, black symbols corresponds to worms feeding on bacteria at OD_600_=3 with no serotonin added. The dotted vertical lines indicate the last point of the linear segment of *F*(*r*), taken as the threshold rate for fast pumping. (B) Zoom-in on the low-rate end of **Fig. 3A** demonstrates the presence of long pauses at all food concentrations. (C) A representative time series of serotonin-induced pumping, illustrating the suppression of very long pauses. (D) Average pumping rates are plotted against the fraction of fast pumping at different concentrations of serotonin in bacteria-free media. (E) *F*(*r*) for wild-type worms and serotonin receptor mutants in bacteria-free media with 5mM serotonin. Zoom-in on the low-rate end demonstrates the suppression of long pauses. (F) Same as in panel (E). Dashed lines are the best-fit lines between 100min^−1^ and 250min^-1^. Dotted vertical lines as in (A).

**Supplementary Movie 1. Time lapse imaging of pharyngeal pumping in the microfluidic device**. Images are taken at a rate of 50 frames per second, shown in real time.

**Supplementary Movie 2. Motion of the posterior bulb during pumping**. The clear contrast between the grinder and its surrounding environment facilitates automatic detection by subtracting consecutive images (red and blue, as in **Fig. 1B**).

